# MOSAIC: a highly efficient, one-step recombineering approach to plasmid editing and diversification

**DOI:** 10.1101/2024.03.22.586135

**Authors:** Marijn van den Brink, Timotheus Y. Althuis, Christophe Danelon, Nico J. Claassens

## Abstract

The editing of plasmids and construction of plasmid libraries is paramount to the engineering of desired functionalities in synthetic biology. Typically, plasmids with targeted mutations are produced through time– and resource-consuming DNA amplification and/or cloning steps. In this study, we establish MOSAIC, a highly efficient protocol for the editing of plasmids and generation of combinatorial plasmid libraries. This quick protocol employs the efficient single-stranded DNA annealing protein (SSAP) CspRecT to incorporate (libraries of) DNA oligos harboring the desired mutations into a target plasmid in *E. coli.* In addition to up to 90% single-target plasmid editing efficiency, we demonstrate that MOSAIC enables the generation of a combinatorial plasmid library spanning four different target regions on a plasmid, in a single transformation. Lastly, we integrated a user-friendly validation pipeline using Nanopore sequencing reads, requiring minimal computational experience. We anticipate that MOSAIC will provide researchers with a simple, rapid and resource-effective method to edit plasmids or generate large, diverse plasmid libraries for a wide range of *in vivo* or *in vitro* applications in molecular and synthetic biology.

GRAPHICAL ABSTRACT

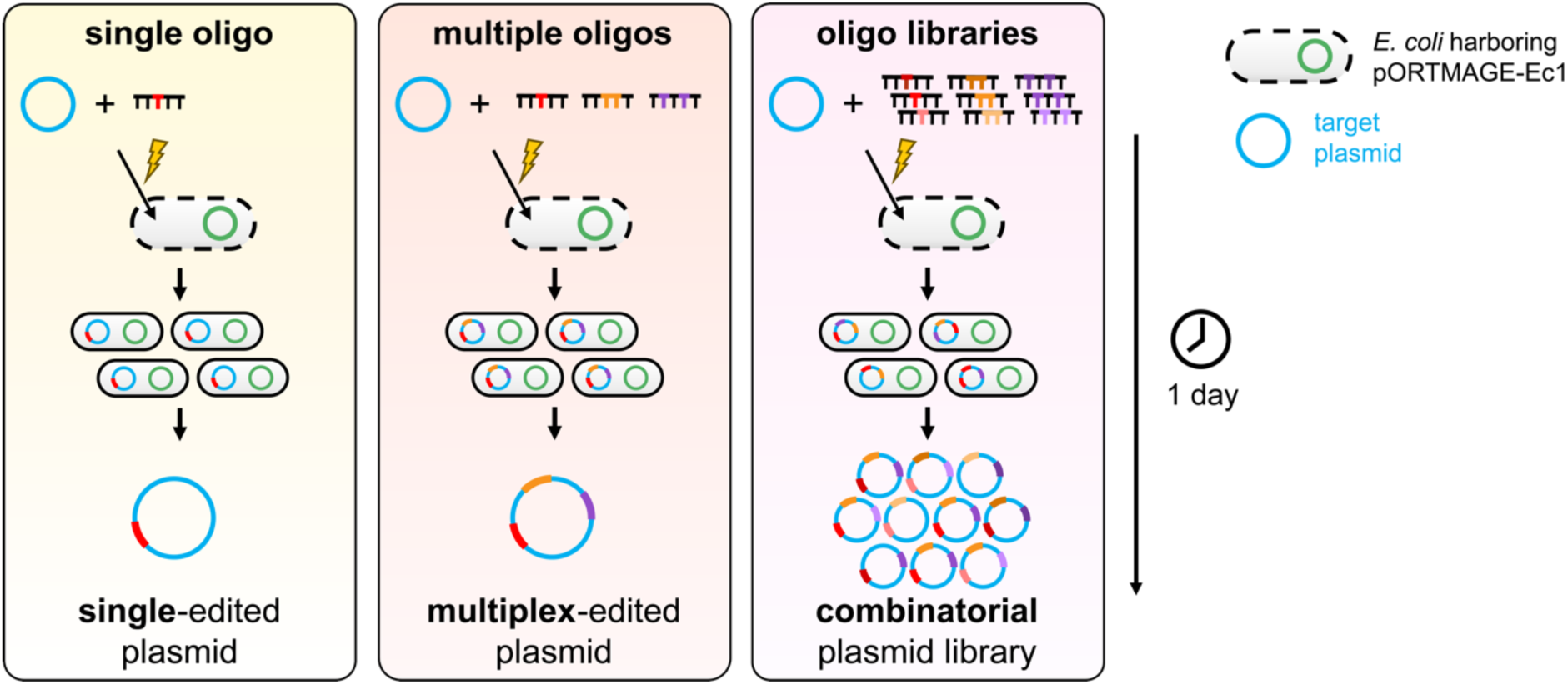

## INTRODUCTION

The development of tools that enable the construction of DNA parts and variants thereof are driving the field of synthetic biology. With the continuing advances in modelling and machine-learning, our predictive capabilities and *a priori* design of functional genetic systems and proteins are rapidly improving (1–5). Still, *in silico* designed genetic parts and proteins often do not behave as expected in the complex genetic and molecular contexts of cells or cell-free expression systems. As a result, efforts in pathway, genetic circuit and protein engineering often benefit from the exploration of wide solution spaces provided by (semi-)rational design and/or computational tools (6, 7).

Plasmid libraries offer an efficient means to test a range of designs *in vivo* or *in vitro*. Generally, mutant plasmids are constructed from DNA fragments amplified by polymerase chain reaction (PCR) with degenerate primers to introduce the desired variation at specific locations. Subsequently, the produced DNA fragments are (re)assembled into a plasmid *in vitro* or *in vivo* by enzymatic assembly (8, 9). For example, site-directed mutagenesis methods based on high-fidelity polymerases and mutagenic primers are widely used (e.g., QuikChange or Q5 site-directed mutagenesis). However, these methods can only diversify one region at a time. Alternative approaches, including restriction enzyme-based and homology-based assembly methods (e.g., Golden Gate, Gibson Assembly, In-Fusion Cloning or Ligase Cycling Reaction) can create plasmids from multiple DNA parts harboring mutations, but are limited in their efficiency and flexibility for the generation of large combinatorial libraries (10–13). Specifically, the number of correctly assembled clones decreases as the number of assembly parts increases. Therefore, researchers need to upscale their experimental efforts or rely on laboratory automation to obtain sufficient numbers of clones to generate larger libraries. These challenges are exacerbated for combinatorial libraries with multiple target sites (multiplexing), large plasmids resulting in low transformation efficiencies, or plasmids containing repetitive DNA. Altogether, there is a need for efficient and flexible strategies to generate large, multiplex plasmid libraries.

Recombineering is a widely used tool to introduce targeted and scarless modifications in bacterial genomes (14). This approach relies on single-stranded DNA (ssDNA) or double-stranded DNA (dsDNA) fragments containing desired mutations, which are introduced into the organism, usually by electroporation. These DNA fragments are then incorporated into replicating chromosomes using phage-derived ssDNA-annealing proteins (SSAPs). This system is extensively employed for genome engineering, especially in *Escherichia coli*, to make large insertions and deletions using dsDNA recombineering and small edits using ssDNA (15, 16). Multiplex automated genome engineering (MAGE) builds on the latter by introducing mutations to many genomic loci at the same time using iterative, automated or manual, editing cycles (17).

Whilst underutilized, ssDNA-mediated recombineering has also been employed to modify plasmids in *E. coli* (18–24). Initially, plasmid recombineering efficiencies of 5-10% were observed for single point mutations with the phage λ-derived SSAP Recβ and two sequential transformations of *E. coli,* first with the target plasmid and then with the mutagenic ssDNA oligos (19). Later, co-electroporation of an optimized ratio of mutagenic ssDNA and the target plasmid yielded editing efficiencies of 20-30% (20). This was further improved to 60% when combined with a co-selection strategy, in which a restriction site on the plasmid is simultaneously mutated, whereafter unmodified variants are eliminated by restriction digestion. Higher efficiencies have also been obtained by combining recombineering with counterselection of non-mutated variants by a CRISPR-Cas nuclease (22, 23). However, this approach requires additional, time-consuming cloning steps and complicates the experimental design and setup. To the best of our knowledge, only a few studies have applied recombineering-based approaches to produce diversified and multiplex plasmid libraries (20, 21, 24). Presumably, the low efficiency of plasmid recombineering and laborious methods relying on co-or counterselection explain the limited application range thus far.

Recently, a systematic screen of phage SSAPs in *E. coli* identified CspRecT, which has a two-fold higher genomic recombineering efficiency than the commonly used Recβ (25). This prompted us to develop MOSAIC: a <u>m</u>ultiplex <u>o</u>ne-<u>s</u>tep SS<u>A</u>P-mediated plasmid diversifi<u>c</u>ation protocol. Its name is derived from *mosaicism*, a phenomenon where mutations give rise to distinct genetic compositions within an organism or a cell population. In this study, we show that MOSAIC’s high plasmid editing efficiency enables the generation of large combinatorial plasmid libraries in a single transformation. Furthermore, MOSAIC employs a validation methodology based on Nanopore long-read sequencing, which quantifies the frequency of (multiplex) library variants directly from the plasmid library sample. We believe that the easy experimental and sequence validation protocols of MOSAIC will facilitate plasmid diversification and expand its range of applications throughout many laboratories.

## MATERIAL AND METHODS

### Reagents and equipment

Chemicals were purchased from Sigma-Aldrich, unless stated otherwise. *m*-toluic acid was dissolved in ethanol at a concentration of 1 M and stored at –20 °C. Plasmids were isolated from bacterial cells using PureYield Plasmid Miniprep System (Promega) or QIAprep Spin Miniprep Kit (Qiagen). Linear DNA was purified using the QIAquick PCR & Gel Cleanup Kit (Qiagen). DNA concentrations were measured using NanoDrop 2000 spectrophotometer (Thermo Fisher Scientific), DS-11 FX spectrophotometer (DeNovix) or Qubit 4 Fluorometer (Invitrogen). Electroporation was performed with 1-mm gap Gene Pulser/MicroPulser electroporation cuvettes (Bio-Rad Laboratories). Different electroporators were used, which all performed robustly for the MOSAIC protocol: the Eppendorf Eporator electroporator (Eppendorf) (1.8 kV) and the ECM 630B electroporator (BTX) (1.8 kV, 200 Ω, 25 µF).

### Strains, cultivation and plasmid construction

Bacterial strains for transformation experiments included *E. coli* K-12 MG1655 (Leibniz Institute DSMZ, Germany), *E. coli* K-12 DH5α and NEB 10-beta *E. coli* (NEB, C3020K). The bacteria were grown in Lysogeny Broth (LB) medium or on LB agar plates containing antibiotics (kanamycin, ampicillin, apramycin) at a concentration of 50 µg/mL unless indicated otherwise. The plasmids used in this study are listed in **Table 1**. pORTMAGE-Ec1 was a gift from the George Church lab (Addgene plasmid #138474; http://n2t.net/addgene:138474; RRID: Addgene_138474). pUC19 was acquired from New England Biolabs. pSEVAb plasmids were cloned according to the method reported earlier (26). Plasmid G555 was constructed by subcloning of the construct containing genes *plsB*, *plsC*, *cdsA* and *pssA* (amplified by primers 1285 ChD and 1286 ChD from plasmid G363) into the backbone of plasmid G340 (amplified by primers 1287 ChD and 1288 ChD) via restriction enzyme digestion (NcoI/XhoI) and ligation. Primers 1285 ChD-1288 ChD are listed in **Table S1**. G340 was constructed as described elsewhere (27). G363 was assembled using a stepwise Golden Gate ligation of six PCR fragments containing independent transcriptional cassettes. First, *plsB*, *plsC* (fragment 1) and *cdsA*, *pssA* (fragment 2) and *tp*, *dnap*, Phi29 origins (fragment 3) were ligated. Then, these three fragments and the pTU1 backbone (Addgene #72934) were ligated to form G363.

**Table 1.** List of plasmids used in this study.

| Plasmid name | Addgene number | Origin of replication | Antibiotic resistance marker | Target region in plasmid for MOSAIC |
| --- | --- | --- | --- | --- |
| pORTMAGE-Ec1 | #138474 | RSF1010 | kanamycin | - |
| pUC19 | #50005 | pUC | ampicillin | *lacZ* |
| pSEVAb827 | #217500 | RK2 | apramycin | *sfGFP* |
| pSEVAb837 | #217501 | pBBR1 | apramycin | *sfGFP* |
| pSEVAb847 | #217502 | pRO1600/ColE1 | apramycin | *sfGFP* |
| pSEVAb867 | #217503 | p15A | apramycin | *sfGFP* |
| pSEVAb887 | #217504 | pUC | apramycin | *sfGFP* |
| pSEVAb897 | #217505 | pBR322/ROP | apramycin | *sfGFP* |
| G555 | #216483 | pUC | ampicillin | RBSs of *plsB*, *plsC*, *cdsA* and *pssA* |

### Recombineering oligos and library design

Mutagenic ssDNA oligos of 89-91 nucleotides were designed to anneal with at least 30 nucleotides at both ends to the target DNA. The ssDNA oligos are listed in **Table S1**. The oligos were modified with two phosphorothioate bonds at the 5’ end. The ssDNA oligos were synthesized and purified by desalting by Sigma-Aldrich (oligos BG31272 and BG31273) or synthesized and purified by HPLC by ELLA Biotech GmbH (Germany) (all other oligos). The oligos were diluted in Milli-Q water to a concentration of 100 µM and stored at –20 °C.

RBS variants were designed using the RBS Library Calculator in the “Optimize Expression Levels” mode (https://salislab.net/software/design_rbs_library_calculator) with the following input parameters: the host organism was *Escherichia coli*; target minimum and maximum translation initiation rates were 1 and 1,000,000, respectively; the genomic RBS sequence was the mRNA sequence from the 5’ end until the start codon (1).

### Plasmid recombineering with the MOSAIC protocol

*E. coli* cells harboring pORTMAGE-Ec1 were streaked from glycerol stocks on a kanamycin-supplemented LB agar plate and grown overnight at 37 °C. The day before the MOSAIC experiment, an individual colony was picked and grown overnight in LB medium supplemented with kanamycin in a shaking incubator at 37 °C and 180-250 rpm. The following day, the overnight culture was diluted 1:100 in LB supplemented with kanamycin in a 50-mL falcon tube and incubated at 37 °C and 180-250 rpm. At an OD_600_ of 0.2-0.3, expression of the pORTMAGE-Ec1 machinery was induced by adding *m*-toluic acid to the culture to a final concentration of 1 mM. Following induction, the cells were incubated for an additional 45 minutes before being placed on ice for 1 hour. To make the cells electrocompetent, the culture was pelleted by centrifugation at 3200 rcf and 4 °C for 10 minutes. Next, the supernatant was carefully decanted before the cells were resuspended in 1 mL of ice-cold Milli-Q water containing 10% glycerol (v/v) and transferred to a 1.5-mL Eppendorf tube. The cells were washed another two to three times. Following the last wash step, the cells were resuspended in 250 μL of ice-cold Milli-Q water per 10 mL of initial culture. Next, 40 µL of cell suspension, 1 ng of target plasmid and 1 μL of 100 μM ssDNA oligos were combined in a 1.5-mL Eppendorf tube. For multi-target MOSAIC reactions, the oligos of interest were premixed at equimolar concentrations and added to the cells to a final concentration of 2.5 µM per oligo or degenerate set of oligos. For the RBS library, degenerate oligos were mixed with one or two additional single oligos per target locus as additional library variants (**Table S1**). Next, 40 µL of the cell-DNA mixture were transferred to a 1-mm gap electroporation cuvette and electroporated. Immediately after electroporation, 960 μL of prewarmed LB were added to the cell suspension and the cells were allowed to recover for 1 hour at 37 °C and 180-250 rpm. Following recovery, single-target transformants were transferred to a 50-mL falcon tube, supplemented with 4 mL of LB containing the appropriate antibiotic and incubated overnight at 37 °C and 180-250 rpm. The next day, the plasmids were isolated from the cells. For multi-target MOSAIC transformations, the recovered cells were plated on large (15 cm diameter) selective agar plates and incubated at 37 °C overnight. The following day, the colonies were counted by hand, whereafter the colonies were scraped off the plate for plasmid isolation. All plasmids were eluted from the plasmid purification columns using Milli-Q water. The DNA purity and concentration were validated by spectrophotometry and fluorometry.

To quantify the number of DNA variants present in single colonies for multi-target MOSAIC with degenerate oligos, six single colonies were picked and grown overnight in ampicillin-supplemented LB (100 µg/mL ampicillin) for plasmid isolation and subsequent Nanopore sequencing.

### Genomic recombineering control experiment

When recombineering was performed on the *E. coli* genome, electroporation of the cells was followed by 1 hour of incubation in 1 mL of LB and, subsequently, 2 hours of incubation in 6 mL of kanamycin-supplemented medium, whereafter the cells were plated on LB agar plates containing kanamycin, 100 µM isopropyl-β-D-thiogalactopyranoside (IPTG) and 100 µg/mL 5-bromo-4-chloro-3-indolyl-beta-D-galacto-pyranoside (X-gal) (Thermo Fisher Scientific). After incubation overnight at 37 °C, the fraction of white colonies relative to the total number of colonies was counted and used as a measure for the genomic recombineering efficiency.

### Retransformation of MOSAIC plasmid mixtures

To separate mutated pUC19 from wild-type pUC19 and pORTMAGE-Ec1 after recombineering, we retransformed competent *E. coli* cells with the resulting plasmid mixtures and screened single colonies for the desired mutations. More specifically, we transformed chemically competent DH5α with 2 ng of the plasmid mixture by heat shock at 42 °C for 45 seconds. After 1 hour recovery in 1 mL of prewarmed LB, 50 µL of cells were plated on prewarmed ampicillin-supplemented (50 µg/mL) agar plates and grown overnight at 37 °C. Single colonies were picked and grown in 3 mL of ampicillin-supplemented (50 µg/mL) LB overnight. The plasmids were isolated from the cells, and the isolation of pure and correctly mutated plasmid was verified by spectrophotometry, gel electrophoresis and Nanopore sequencing.

To separate the RBS plasmid library from pORTMAGE-Ec1, 40 µL of NEB 10-beta electrocompetent *E. coli* cells (C3020K) were transformed with 2.5 µL (155 ng) of plasmid mixture by electroporation, following the instructions from NEB (Electroporation Protocol C3020). All cells were plated on five large (15 cm diameter) ampicillin-supplemented (100 µg/mL) agar plates and grown overnight at 37 °C. The colonies were scraped off the plates for plasmid isolation. The plasmids were then linearized at a unique restriction site outside of the mutagenized region of interest using StuI (0.8 U/µL) in rCutSmart buffer to ensure that the obtained Nanopore sequencing reads spanned the full region of interest. The linearized DNA was purified by excision of the expected band from a 0.7% agarose gel and further purification using the QIAquick PCR & Gel Cleanup Kit (Qiagen). The linear DNA was diluted in Milli-Q water to a concentration of 99 ng/µL and sequenced by Nanopore sequencing.

### Nanopore sequencing and analysis

Nanopore sequencing was performed with the mixture containing target plasmid and pORTMAGE-Ec1 unless indicated otherwise. Samples were prepared by diluting the DNA in Milli-Q water to a concentration of 30-40 ng/µL as quantified by Qubit. Nanopore sequencing was performed by Plasmidsaurus (Oregon, US). To extract the sequencing reads that map to the target plasmid, the reads were filtered based on size (the target plasmid size plus and minus 100 bp) and mapped to the wild-type DNA sequence of the target plasmid using the *Filter FASTQ reads by quality score and length* (28) and *Map with minimap2* (29) tools, respectively (accessed in the Galaxy web platform (https://usegalaxy.org)) (30). To minimize the numbers of insertions/deletions rather than mismatches during mapping, the minimap2 alignment parameters gap open penalty for deletions and insertions were increased from 4 (default) to 16 and from 24 (default) to 48, respectively, for the MOSAIC experiments incorporating 18 nt-insertions and deletions in pUC19 and diversifying the four RBSs in plasmid G555. For the 18-nt mismatch samples, these were increased to 32 and 72, respectively. For the 18-nt insertion samples, the reads were mapped to the designed modified DNA sequence instead of the wild-type sequence. If the reads from multiple Nanopore sequencing runs of the same sample were used for analysis, the FASTQ datasets were first merged using *Concatenate datasets tail-to-head* tool in the Galaxy web platform.

In-house developed R scripts were run in Rstudio (Version 1.1.456) to determine the editing efficiency (available at https://dx.doi.org/10.4121/4464ab86-9214-49b3-a808-10ca655385a6). The sequential steps in the analysis pipeline are illustrated in **Figure S1**. In short, the DNA sequences from the target loci were extracted from the mapped reads and filtered based on the per-base quality scores recorded in the FASTQ files; the target sequences that contained at least one base with a score lower than 50 were excluded from the analysis. Then, the target sequences were identified as the wild type or as successfully mutated based on 100% similarity. The fraction of mutated sequences relative to the total number of target sequences was used to determine the editing efficiency. If the plasmid was modified in multiple loci, the number of mutated target loci was also counted per plasmid. To that end, an additional filtering step was applied to remove all reads that did not span the full sequence from the first to the last target region.

### Statistics

The editing efficiencies described in the main text were averaged over at least three biological replicates. The mean and standard deviation are given where indicated.

## RESULTS AND DISCUSSION

### CspRecT-mediated recombineering achieves ∼85% single-locus plasmid editing efficiency

We investigated the efficiency of plasmid recombineering using SSAP CspRecT expressed from plasmid pORTMAGE-Ec1 (**Figure 1A**) (25). The plasmid recombineering efficiency was first tested with the high-copy number plasmid pUC19. Two types of ssDNA oligos were designed to introduce a single-nucleotide deletion or a three-nucleotide mismatch in the *lacZ* gene on pUC19. The oligos contained two phosphorothioate bonds at the 5’ ends to protect against degradation by exonucleases *in vivo* (17). Because pUC19 replicates unidirectionally in a DNA sequence-controlled manner (31–33), we designed the oligo sequences such that they target the lagging strand during plasmid replication, as this is believed to lead to the highest efficiency during recombineering (14, 20, 34). The ssDNA oligos (2.5 µM) were co-electroporated with 1 ng of pUC19 plasmid into electrocompetent *E. coli* MG1655 expressing CspRecT and the dominant negative *E. coli* MutL mutant (EcMutL^E32K^) for temporal repression of mismatch repair. Usually, deleterious mutations in *lacZ* can be quantified using blue-white screening on an LB agar plate with X-gal. However, as plasmid recombineering leads to mixed plasmid populations in single colonies, we determined the editing efficiencies by DNA sequencing. Hence, after overnight growth, the plasmids were isolated, and the editing efficiency was quantified from Nanopore sequencing reads. The Nanopore sequencing analysis pipeline is shown in **Figure S1**. Only reads with a quality threshold ≥50 in the target region were used to reduce the chance of incorrect detection of mutations to < 1% (**Figure S2**).

**Figure 1.**
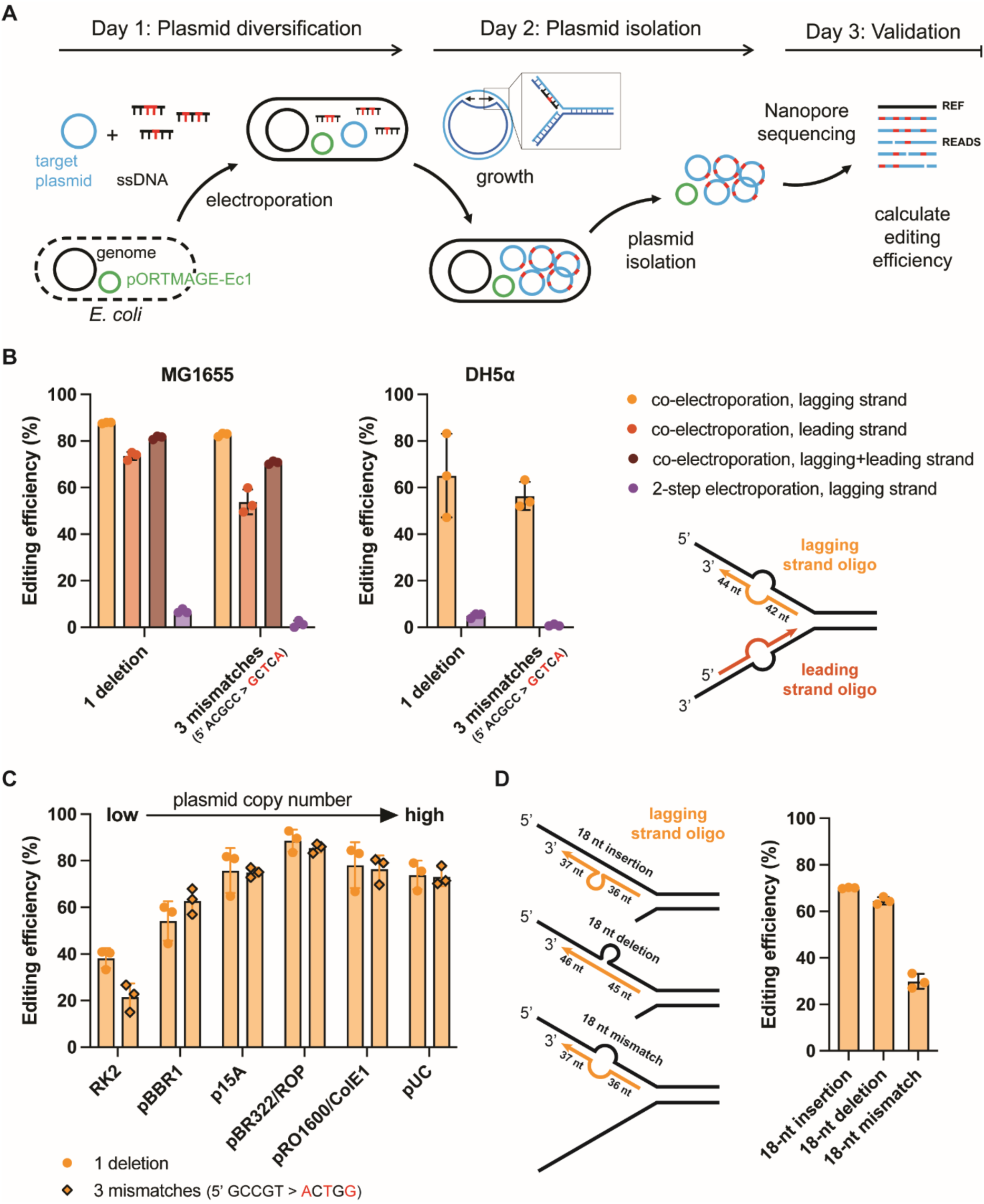
Plasmid editing efficiency of MOSAIC. **A)** Schematic representation of MOSAIC’s protocol for plasmid editing using ssDNA recombineering. The expression of SSAP CspRecT and the dominant negative mutant EcMutL^E32K^ is induced in *E. coli* cells harboring pORTMAGE-Ec1. The cells are made electrocompetent and transformed with the target plasmid and mutagenic ssDNA oligos. During plasmid replication, the oligos anneal to one of the two DNA strands in the replication fork with the help of CspRecT, introducing mutations into the plasmid sequence. Plasmids are then isolated from the cells, and the editing efficiency is calculated from Nanopore sequencing reads. **B)** Plasmid editing efficiencies for the incorporation of a single-nucleotide deletion or a three-nucleotide mismatch in the *lacZ* gene in high-copy number plasmid pUC19. We investigated the effects of lagging strand versus leading strand oligos, the use of co-or 2-step electroporation and the use of *E. coli* strains MG1655 or DH5α. **C)** Plasmid editing efficiency for the incorporation of a single-nucleotide deletion or a three-nucleotide mismatch in the gene encoding sfGFP on the pSEVAb plasmids with different origins of replication. **D)** DNA editing efficiency for the incorporation of an 18-nt long insertion, deletion or mismatch in the *lacZ* gene on the high-copy number plasmid pUC19. For panels C and D, lagging strand oligos were used, and they were co-electroporated into *E. coli* MG1655 together with the plasmid DNA.

Remarkably, the editing efficiency after a single round of MOSAIC was 88% and 83% for the single-nucleotide deletion and three-nucleotide mismatch, respectively (**Figure 1B**). As expected, the use of complementary oligos that bind the leading strand in the replication fork resulted in lower editing efficiencies (74% and 54% for the single-nucleotide deletion and three-nucleotide mismatch, respectively) (**Figure 1B**). The addition of both the leading and lagging strand-targeting oligos did not further improve the editing efficiency. The observed editing efficiencies were more than double those previously reported for plasmid editing with the Redβ SSAP (20). This coincides with the previously observed two-fold increase in genomic editing efficiency with CspRecT versus Redβ (25), highlighting the large impact of SSAP on the editing efficiency for both genomic and plasmid recombineering.

Importantly, when the oligos were electroporated into bacteria already harboring the target plasmid (i.e., “2-step electroporation”), the fraction of edited plasmids was very low (<7%) (**Figure 1B**). So, co-electroporation of the target plasmid and ssDNA is key to reach high plasmid editing efficiencies. This is in agreement with a previous study using Redβ for plasmid recombineering (20). The large effect of co-electroporation is likely explained by the fact that recombineering is most effective during plasmid replication. When a single plasmid or a low number of plasmids enters the cell, the plasmid(s) will likely be rapidly replicated many times to reach the copy number at which the plasmid is maintained in the cells.

Overall, plasmid recombineering resulted in much higher efficiencies than recombineering on the *E. coli* genome, whose highest reported efficiency is ∼50% but in our hands reached only 14% based on a blue-white screening (**Figure S3A**) (25). We also tested the MOSAIC protocol in *E. coli* DH5α, which is routinely used for transformation and cloning purposes. However, the observed plasmid editing efficiency in this strain was lower than in *E. coli* MG1655 (**Figure 1B**). Hence, unless stated otherwise, *E. coli* MG1655 was used for recombineering in the remainder of this study.

### Higher-copy plasmids are edited more efficiently than low-copy plasmids

As we hypothesized that the high editing efficiency was coupled to a high plasmid replication rate, we anticipated that higher-copy plasmids with comparatively higher replication rates after electroporation would be edited more efficiently than lower-copy plasmids (20). To test this, we applied MOSAIC to a series of pSEVAb vectors that differed only in their origins of replication and, consequently, the copy number at which they are maintained in *E. coli* (26). The selected origins of replications were RK2 (low-copy number), pBBR1, p15A, and pBR322/ROP (medium-copy number), and pRO1600/ColE1 and pUC (high-copy number) (35, 36) (**Figure S4**). We designed oligos to incorporate a deletion or a three-nucleotide mismatch in the gene encoding sfGFP present in all plasmids. We identified the plasmid leading and lagging strands based on the known class B theta replication mechanism of ColE1 and ColE1-like origins (pRO1600/ColE1, pUC, pBR322 and p15A) (31–33, 37, 38), and similarly for the class A theta replication mechanism of the RK2-plasmid origin (33, 39, 40). Based on this, we tested the oligos targeting the lagging strand assuming this would lead to the highest recombineering efficiency. To the best of our knowledge, the precise replication mechanism of pBBR1-derived plasmids is still unknown. Therefore, we tested both (reverse complementary) oligos for this plasmid, which performed equally well (**Figure S3B**). As anticipated, the vector with the low-copy RK2 origin of replication was edited with the lowest efficiency, on average 30%, followed by 58% for the pBBR1 origin of replication (**Figure 1C**). Surprisingly, the four other plasmids that we evaluated were all modified with 70-90% efficiency. As such, it appears that beyond a certain copy number, additional replication events no longer increase the efficiency at which a single oligo is incorporated. Such a threshold might be due to a saturation effect, possibly caused by exceeding a time window during which the recombineering machinery and/or oligos are sufficiently active, and may represent an upper boundary for MOSAIC.

### Large insertions and deletions are incorporated with high efficiency

To probe the potential broad applicability of MOSAIC, we investigated if a high editing efficiency could still be obtained with a substantially larger number of mutations per oligo. If so, larger regions, such as regulatory sequences (e.g., promoters, RBSs and operator sites), could readily be inserted, deleted, replaced or diversified. Hence, we designed three ssDNA oligos that incorporate an 18-nt wide insertion, deletion or mismatch into the *lacZ* gene on pUC19. The efficiency for the insertion and deletion was 65-70% (**Figure 1D**). The efficiency for substituting 18 nucleotides was lower (30%), but still sufficient for many of the aforementioned applications. Additionally, the large substitution was 5-fold more efficient than the earlier reported substitution of identical length in the *E. coli* genome (25). Altogether, these results demonstrate that MOSAIC is a powerful method to edit and diversify both small and larger regions of plasmid DNA.

### One round of MOSAIC yields a large multiplex plasmid library

Next, we investigated if we could apply MOSAIC to create a combinatorial plasmid library of ∼10^4^ variants in a single electroporation step. As a proof-of-principle, we diversified the ribosome binding sites (RBSs) of four genes encoding a phospholipid synthesis pathway on the pUC19-derived plasmid G555 (**Figure 2A**) (41). Mutations in RBS sequences are expected to change the absolute and relative abundances of the four encoded proteins, hence phospholipid production. This enzymatic cascade from *E. coli* has been reconstituted in cell-free systems (41) and is amenable to phenotypic characterization of RBS modifications both *in vivo* and *in vitro*. To modulate translation, RBS Calculator was used to design RBS variants with a wide range of predicted translation initiation rates (1). The resulting variants contained up to 7 mismatches per RBS relative to the wild-type DNA and yielded a total library of 13 x 11 x 9 x 9 = 11,583 theoretical DNA variants (**Figure 2B**). Because two of the four target sites were identical, three degenerate oligo libraries targeting four different sites on the plasmid were sufficient to produce the combinatorial library. In a single electroporation reaction, the G555 plasmid and a 1:1:1 mix of the three degenerate oligo libraries were transformed into *E. coli* MG1655 cells expressing CspRecT and EcMutL^E32K^. G555 contains multiple highly similar sequences (e.g., transcriptional promoters, RBSs and terminators) and is prone to recombination in *E. coli* MG1655. As such, the cells were directly plated on selective agar plates to prevent that some cells, harboring incomplete plasmids with a lower expression burden, outcompete the cells with full-length plasmids. Such competition would be a potential risk if the libraries were cultivated in liquid media. Following incubation overnight, the plasmid libraries from four plasmid recombineering transformations were isolated and sequenced.

**Figure 2.**
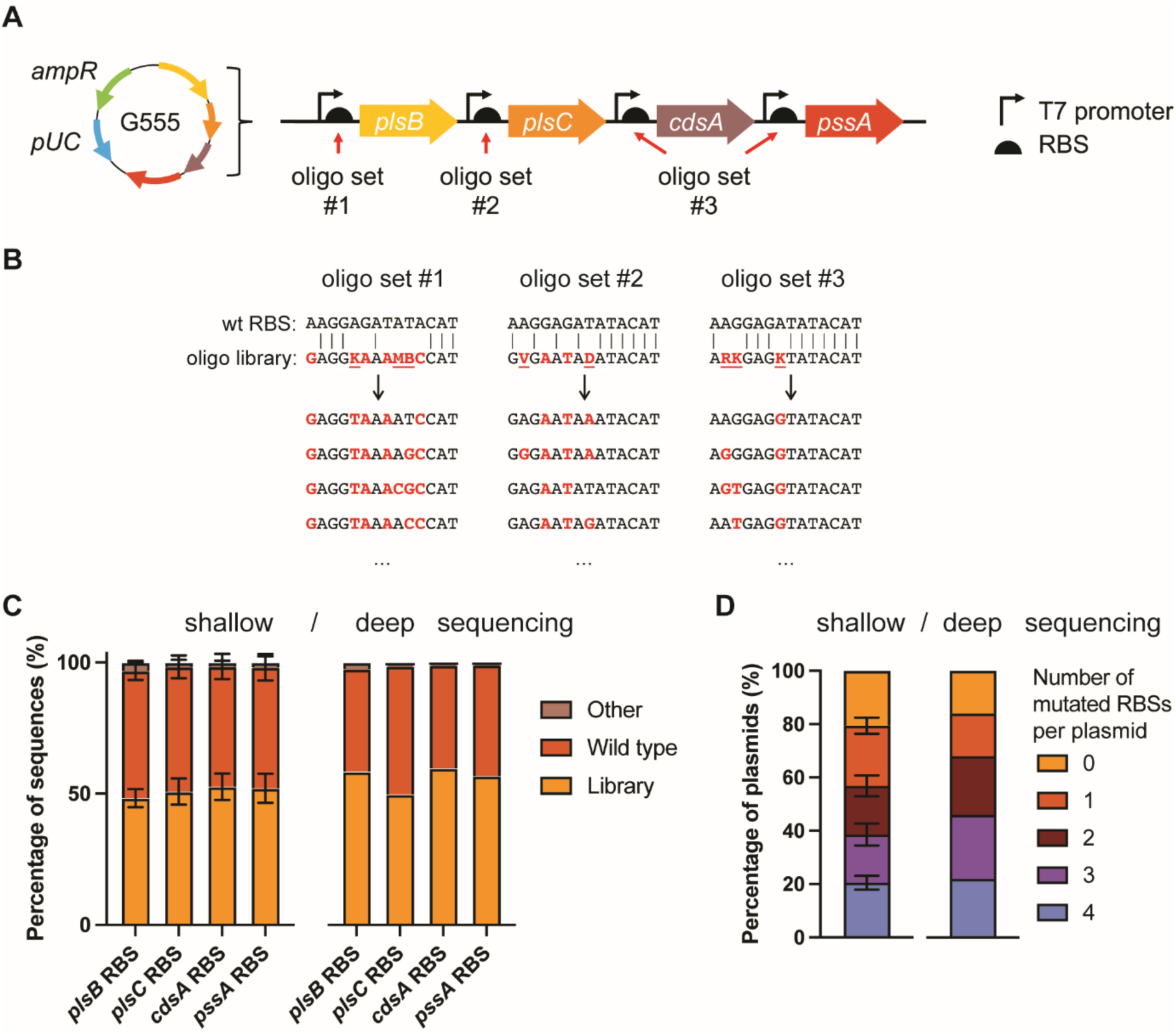
Construction of a multiplex plasmid library with four diversified RBSs using MOSAIC. **A)** pUC-derived target plasmid G555 containing four genes from the Kennedy phospholipid biosynthesis pathway (*plsB*, *plsC*, *cdsA* and *pssA*). Three ssDNA oligo sets of degenerate sequences (9-13 variants per set) targeted four regions on plasmid G555. **B)** The three sets of ssDNA oligos were designed using RBS Calculator (1) to target the RBSs of four genes. Letters in red indicate changes relative to the wild-type DNA. Underlined letters indicate degenerate nucleotides. **C)** Percentage of wild-type and library sequences per target locus, quantified from shallow Nanopore sequencing (left graph, n=4 with each 150-300 reads per target locus) or a deep sequencing dataset (right graph, n=1 with 279,454 reads per target locus). **D)** Number of mutated RBSs per plasmid, quantified from shallow Nanopore sequencing (left graph, n=4 with each 20-50 full-length reads) or the deep sequencing dataset (right graph, n=1 with 279,454 full-length reads).

Through Nanopore sequencing, the target loci are (physically) linked in a single read enabling the identification of the full genotype of each plasmid variant. Using the analysis pipeline outlined in **Figure S1**, the RBS variants in the Nanopore reads mapping to the phospholipid synthesis pathway genes were compared to the library variants designed by RBS Calculator and to the wild-type RBSs, and their frequencies were counted. Based on 150-300 Nanopore reads, we calculated the per-locus editing efficiencies as 48% ± 3% (*plsB* RBS), 51% ± 5% (*plsC* RBS), 53% ± 5% (*cdsA* RBS) and 52% ± 6% (*pssA* RBS) (mean ± standard deviation, n=4) (**Figure 2C**, left graph). Thus, the editing frequency was consistent across the target sites and across replicates. With an average editing frequency of 51%, we expected that approximately 7% (i.e., 0.51^4^) of the DNA molecules would have mutations in all four RBS sequences. However, we observed that 21 ± 3% (n=4) of the sequenced library variants had all four RBS sequences mutated (**Figure 2D**, left graph). This represents an unexpectedly high multiplexing efficiency of 21% after a single round of plasmid engineering. This suggests that there is a subpopulation of cells or plasmids with a higher-than-average editing efficiency and, thus, a higher chance that all loci are edited simultaneously. In addition, 79% of the library variants had at least one target site mutated, 57% at least two, and 39% had at least three target sites mutated. In contrast, previous work seeking to modify multiple sites on a plasmid with the Redβ recombinase observed >1 mutation in only 25% of their library (20). Additionally, Higgins *et al*. observed 2 mutated target sites in 25% of their population after 5 rounds of plasmid recombineering (21).

To estimate the library coverage, we sequenced one of the four plasmid libraries in more depth. In the previous, shallow sequencing, we obtained only ∼50 high-quality reads that spanned all target loci in a single read. To obtain as many reads spanning the full mutagenized region of interest on G555 as possible, the library was purified from pORTMAGE-Ec1 by retransformation and linearized at a unique restriction site outside the region of interest. We obtained ∼300,000 high-quality reads spanning all four target sites. Of these reads, ∼60,000 or 22% had all four RBS sequences mutated, echoing the multiplexing efficiency we observed in our shallow sequencing dataset (**Figure 2D**). Furthermore, all designed RBS variants were present with similar frequencies (**Figure S5**), and 99% of the library variants were present in the same order of magnitude (**Figure S6A**), suggesting unbiased RBS diversification. Altogether, we detected 2,839 of the 11,583 designed library variants, representing a library coverage of at least 25%. We believe this value represents a minimum, as 38% of the library variants were represented by one or two reads, suggesting that deeper sequencing may be required for full coverage of the DNA library. This was also indicated by the rarefaction curve (**Figure 6SB**), which was still increasing at 60,000 reads, suggesting that the sequencing depth is limiting the number of variants detected and the library size is likely larger than 2,839 variants.

Notably, only ∼1-3% of RBS sequences were neither the wild-type sequence nor a designed library variant (“Other”, **Figure 2C**). These unintended mutations are likely sequencing errors that were not excluded by our analysis pipeline (**Figure S2B**), mutations incorporated by oligos intended to bind at other target sites, and/or spontaneous mutations retained due to the suppression of the mismatch repair system during recombineering. The low fraction of unintended mutations is particularly noteworthy given that the oligos share an almost identical 40-nt left arm. It appears that having only one unique arm in the oligos is sufficient for the specific introduction of mutations. This is especially useful for mutating, for example, multiple 5’UTR or terminator regions with high sequence similarity.

Lastly, we tested whether performing successive rounds of plasmid recombineering leads to the accumulation of mutations at the four target regions in plasmid G555. As G555 is prone to recombine in MG1655, we performed these experiments in DH5α harboring pORTMAGE-Ec1. Following each round of plasmid recombineering, the target plasmid was selected for in liquid media overnight, the plasmids were isolated, and 30 ng was used as the starting point for the subsequent round. The editing efficiencies were quantified from Nanopore sequencing reads for three successive rounds. Despite the slightly lower plasmid editing efficiency observed for DH5α (**Figure 1B**), we observed an increase in the number of mutated RBSs per plasmid with each subsequent round of plasmid recombineering (**Figure S7A**). Interestingly, the *plsB* locus was edited roughly >2 fold more often than the other loci (**Figure S7B**). Altogether, these results demonstrate that multiple rounds of retransformation and recombineering can be used to increase the frequency of mutations across multiple target loci on a plasmid.

To conclude, these results demonstrate that MOSAIC enables the generation of large, diversified plasmid libraries in a single transformation. Moreover, key parameters for library characterization can be accurately determined by commercial, fast and low-cost Nanopore sequencing.

### Attaining clonality: purifying plasmid mixtures after recombineering

In assembly-based cloning methods, cells or colonies typically harbor a single plasmid variant after transformation. In contrast, plasmid recombineering yields colonies with multiple DNA variants, as mutations are incorporated in the plasmids after uptake by the cells. Single-locus plasmid edits yield colonies harboring both the mutant and wild-type plasmids. To determine the number of plasmid variants present in a single colony following multiplex plasmid recombineering, we miniprepped and sequenced plasmid mixtures from six single colonies following our G555 RBS diversification experiment. On average, we found 6 ± 4 (n=6 colonies) different DNA variants per colony (**Table S2**). To ensure that the modified plasmids are suitable for an application of interest, the following steps can be taken to attain clonality and/or remove pORTMAGE-Ec1 from the plasmid pool (**Figure 3A**).

**Figure 3.**
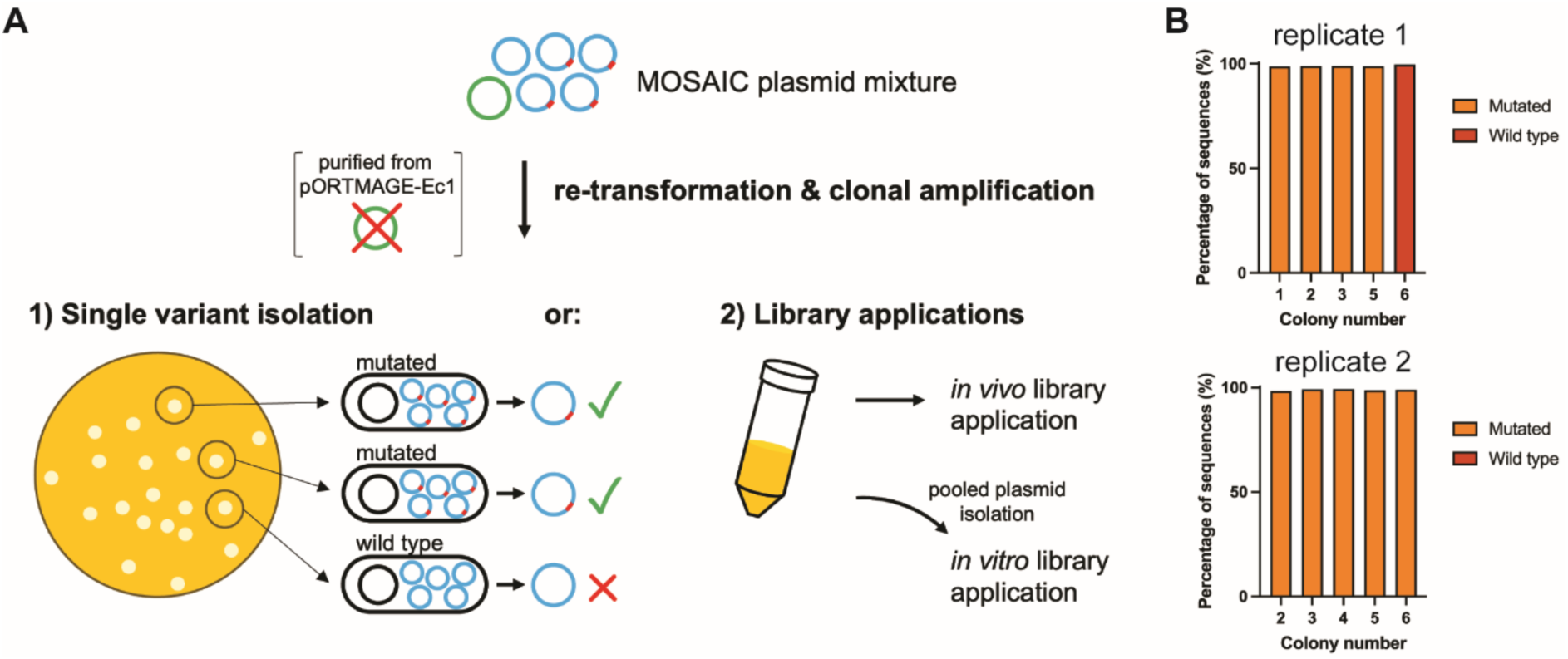
Retransformation of plasmid mixtures generated by plasmid recombineering is sufficient to remove pORTMAGE-Ec1 and establish clonality. **A)** Pure, mutated plasmids can be isolated by single-colony picking after retransformation. Plasmid libraries can be purified from pORTMAGE-Ec1 by retransformation, using either a cloning strain (e.g., DH5α or NEB 10-beta electrocompetent cells) or presumably the strain of interest for an *in vivo* application. For *in vitro* applications, libraries can be isolated from the cloning strain after retransformation. **B)** Clonality of re-transformants from single-edited plasmids.

To obtain clonality and remove the pORTMAGE-Ec1 helper plasmid, plasmids can be isolated from the MOSAIC-derived population and used for retransformation of *E. coli* or other hosts, while only selecting for target plasmids. We demonstrate this by retransforming *E. coli* with the plasmid mixture from an earlier, single-locus pUC19 editing experiment. After transformation and growth of *E. coli* DH5α on plates selective for pUC19, plasmids from several colonies were individually extracted, analyzed on agarose gel and sequenced. Following retransformation, the band in the gel corresponding to the size of pORTMAGE-Ec1 was no longer present, suggesting the successful removal of pORTMAGE-Ec1 (**Figure S8**). This was confirmed by Nanopore sequencing as none of the colonies yielded reads that mapped to pORTMAGE-Ec1. More importantly, the sequencing data showed that each individual colony was associated with a single genotype; there was a strict separation of mutated and wild-type plasmids (**Figure 3B**). Four of the five (replicate 1) and five out of five (replicate 2) colonies harbored the mutated variant, which is in line with the high single-locus editing efficiency of plasmids with this origin of replication (∼83%). With these results, we verify clonality at the sequence level and demonstrate that modified and unmodified plasmids can be untangled after plasmid recombineering by a simple retransformation.

Due to the attained clonality, we presume that libraries generated by plasmid recombineering can be applied directly *in vivo* by retransformation into a strain of interest, as long as a sufficient number of transformants are obtained to ensure adequate library coverage. Alternatively, libraries can be applied *in vitro* following pooled plasmid isolation. To demonstrate recovery of the mutagenized G555 library whilst purifying the mixture of pORTMAGE-Ec1, we retransformed 155 ng of the mutagenized RBS plasmid library into highly electrocompetent NEB 10-beta cells yielding around 500,000 colonies, which is sufficient to cover the full library. Subsequent pooled plasmid isolation, digestion and Nanopore sequencing confirmed complete removal of pORTMAGE-Ec1, as no sequencing reads could be mapped to pORTMAGE-Ec1. Alternatively, the target DNA could be amplified by PCR directly from the mixture of target plasmids and pORTMAGE-Ec1.

## CONCLUSIONS

In contrast to state-of-the-art plasmid editing methods, such as Golden Gate and Gibson assembly, MOSAIC does not require PCR or plasmid assembly from fragments. The only requirements for MOSAIC are fast-to-order mutagenic oligos and a publicly available *E. coli* strain harboring pORTMAGE-Ec1. A simple co-electroporation of the target plasmid and oligos is sufficient to perform the desired mutagenesis. This study has built on earlier work by increasing the complexity of plasmid engineering through the new, more efficient pORTMAGE-Ec1 system. The remarkably high editing efficiency and user-friendly library sequence characterization protocols, made accessible to scientists with minimal computational experience, now enable the use of plasmid recombineering and library sequence analysis on a regular basis.

The simplicity of MOSAIC’s protocol lends itself to a plethora of *in vivo* and cell-free synthetic biology applications ranging from protein engineering to the optimization of natural or synthetic metabolic pathways (13, 42–44). More specifically, MOSAIC could enable the rapid prototyping of pathway or enzyme variants when coupled to high-throughput phenotypic screening or growth-coupled selection approaches to isolate well-performing variants. These isolated plasmid variants could in principle be rapidly subjected to further rounds of diversification using MOSAIC during subsequent design-build-test-learn cycles.

To synthesize large plasmid libraries, a high transformation efficiency is required. In contrast to cloning-based library synthesis methods, plasmid recombineering benefits from the transformation of preassembled plasmids. In other words, transformation efficiency is not limited by incomplete assemblies. However, the presence of full and partial wild-type plasmid variants alongside the mutant DNA variants in combinatorial MOSAIC libraries is currently unavoidable. As such, increasing the number of target sites or the variability at each site may hinder library coverage. The library coverage could be increased by performing multiple reactions in parallel or improving the transformation efficiency. Alternatively, the fraction of fully mutated library variants could be improved by employing iterative rounds of MOSAIC, co-selection using restriction enzymes, or CRISPR-Cas-based counterselection (20, 22). Another promising strategy is to restrain the library size by computational design. Excitingly, recent efforts to preselect library variants from a larger pool using machine learning prior to wet lab characterization showed promising results in generating small but smart libraries to accelerate the evolutionary optimization (45). All in all, we anticipate that MOSAIC, combined with the continuing advances in computationally aided design of genetic and protein libraries, will enable the rapid exploration of biological solution spaces throughout many labs and research projects.

## DATA AVAILABILITY

The data underlying this article are available in the 4TU.ResearchData repository at https://dx.doi.org/10.4121/4464ab86-9214-49b3-a808-10ca655385a6.

## AUTHOR CONTRIBUTIONS

MvdB and TYA contributed equally to this study. Conceptualization and Methodology: MdvB, TYA, CD and NJC. Investigation and formal analysis: MvdB and TYA. Software and Data Curation (analysis pipeline development): MvdB. Visualization and Writing – Original draft: MvdB and TYA. Writing – Review & Editing: MdvB, TYA, CD and NJC. Supervision & Funding acquisition: CD and NJC.

## SUPPORTING INFORMATION

Additional data for Nanopore sequencing, bacterial plating, agarose gel electrophoresis, as well as a list of DNA oligo sequences used in this study.

## ACKNOWLEDGEMENTS

We thank Ana Restrepo Sierra for her contribution to the conceptualization and design of the RBS library. We also thank Suzan Yilmaz for assistance in setting up the recombineering experiments. We are thankful to Akos Nyerges for providing advice on recombineering and the use of pORTMAGE. Lastly, we thank Haroun Bensaadi for performing preliminary recombineering experiments. MvdB, CD and NJC acknowledge financial support from The Netherlands Organization of Scientific Research (NWO/OCW) Gravitation program Building A Synthetic Cell (BaSyC) (024.003.019). TYA and NJC acknowledge funding from an NWO-XL grant (OCENW.XL21.XL21.007). CD acknowledges funding from ANR (ANR-22-CPJ2-0091-01). NJC also acknowledges support from an NWO-Veni Fellowship (VI.Veni.192.156).

