## Supporting Information for "MOSAIC: a highly efficient, one-step recombineering approach to plasmid editing and diversification"

† Joint Authors


**Content:**


Figure S1 | Nanopore sequencing analysis pipeline used in this study.

Figure S2 | Analysis of Nanopore sequencing reads for wild-type pUC19 and wild-type G555 using different quality thresholds.

Figure S3 | Editing efficiency targeting the *E. coli* genome or pSEVA plasmids using lagging or leading strand oligos.

Figure S4 | Ratio of the number of Nanopore sequencing reads for the pSEVAb plasmids to those for pORTMAGE-Ec1.

Figure S5 | Frequency of each of the RBS variants per target locus.

Figure S6 | Sequencing depth for the retransformed RBS library.

Figure S7 | Three successive rounds of MOSAIC increase the RBS library diversity of plasmid G555.

Figure S8 | Isolation of mutated pUC19 by retransformation.

Table S1 | DNA oligos used in this study.

Table S2 | Number of sequenced DNA variants per colony.

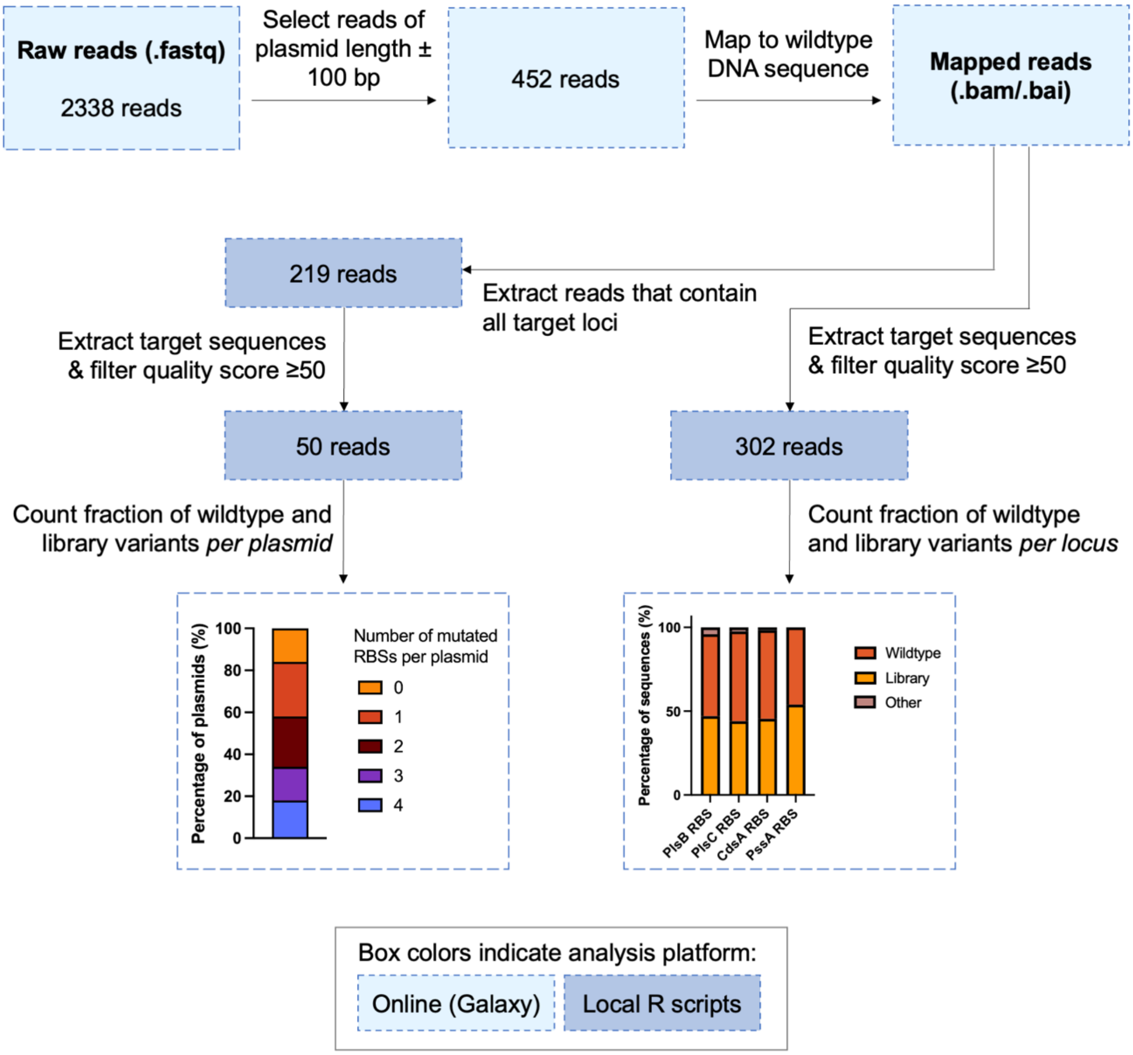


**Figure S1. Nanopore sequencing analysis pipeline used in this study.** As an example, the graphs and numbers of reads are given for a single MOSAIC sample where four RBSs were diversified in four different loci on plasmid G555.

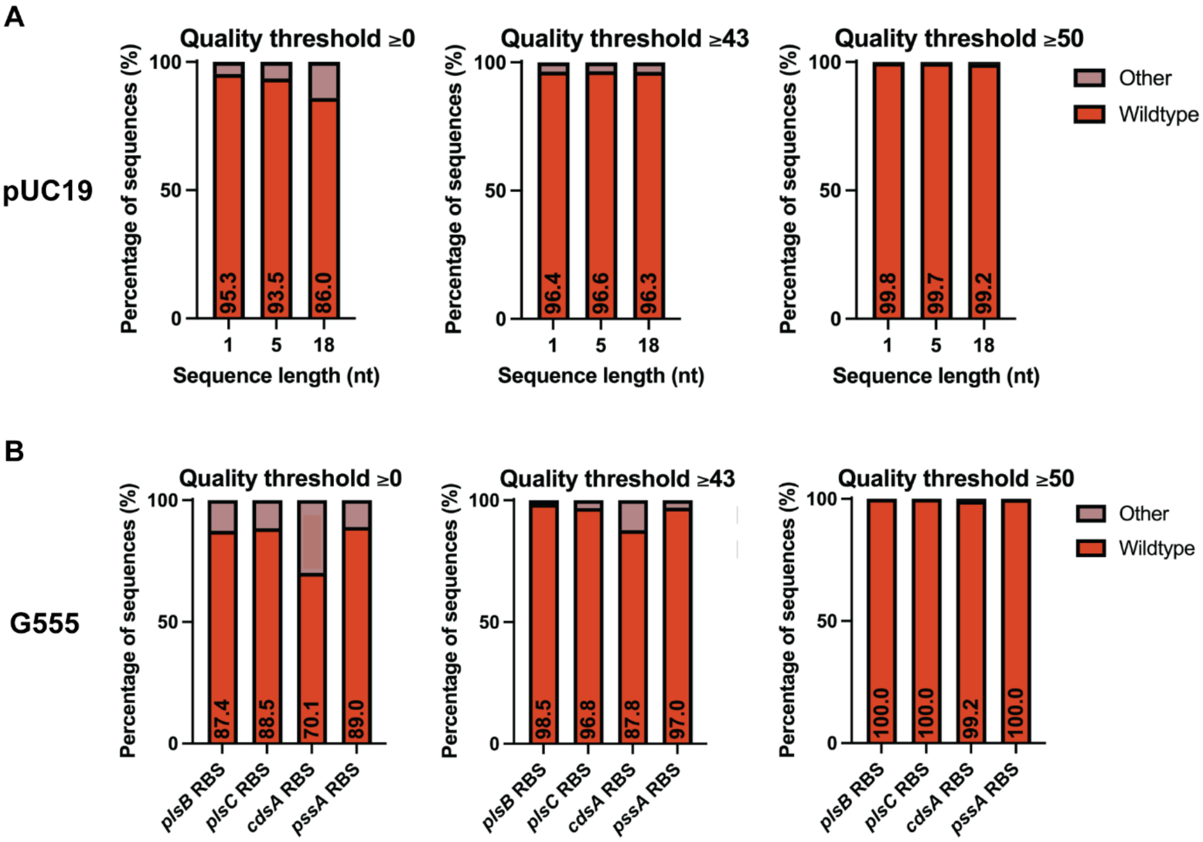


**Figure S2. Analysis of Nanopore sequencing reads for wild-type pUC19 and wild-type G555 using different quality thresholds. A)** Different quality thresholds applied to specific loci in wild-type pUC19 sequencing reads (1, 5 or 18 nt long) that correspond to the target loci in the experiments of Figure 1B and Figure 1D. **B)** Different quality thresholds applied to the specific loci in wild-type plasmid G555 sequencing reads (20 nt long) that correspond to the target loci in the experiments of Figure 2 and Figure S5. While for short target regions of 1-5 nucleotides the fraction of sequencing errors was relatively small without quality control (4-6%), for larger target regions (18-20 nt) the fraction of sequencing errors became large (11-30%). The value 50 was applied to the experiments in this study as the quality score threshold to minimize sequencing errors based on the sequencing data of the wild-type plasmids to <1%, while maintaining as many sequences for analysis as possible.

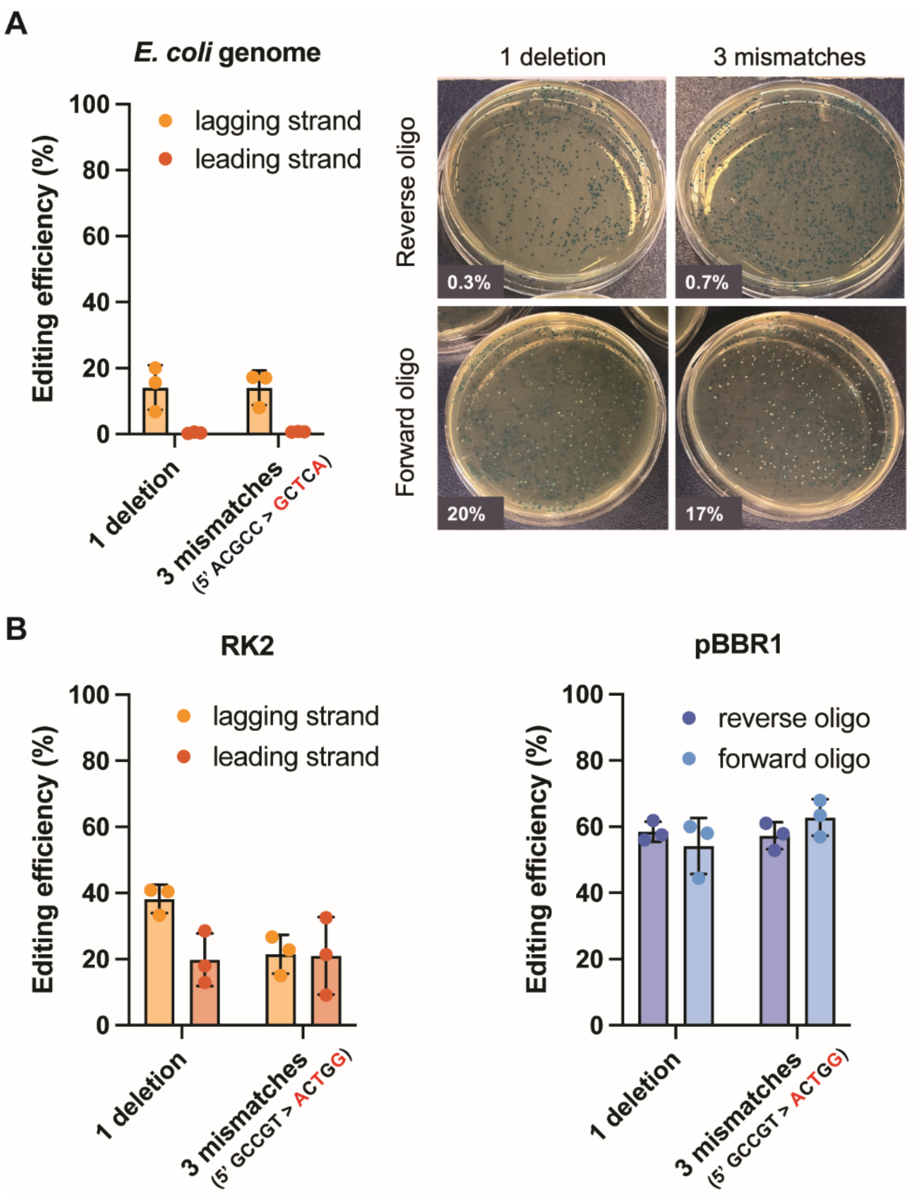


**Figure S3. Editing efficiency targeting the *E. coli* genome or pSEVA plasmids using lagging or leading strand oligos. A)** DNA editing efficiency for the incorporation of one deletion or three nucleotide mismatches in the *lacZ* gene in the genome of *E. coli* MG1655 harboring pORTMAGE-Ec1. The editing efficiencies were quantified by blue-white screening. The deletion in the *lacZ* gene caused a frameshift resulting in white colonies (deficient β-galactosidase), and the three substitutions led to a premature stop codon, resulting in light-blue colonies, while the fully functional variants appeared as dark blue colonies. **B)** DNA editing efficiency for the incorporation of one deletion or three nucleotide mismatches in the *sfGFP* gene on RK2-derived plasmid pSEVAb827 (left) or pBBR1-derived plasmid pSEVAb837 (right). As it is unknown which of the strands are the leading and lagging strands for pBBR1, we indicate the reverse complementary oligos with "forward" and "reverse" oligo, which anneal to the noncoding and coding strand of the *sfGFP* gene, respectively. The oligos were co-electroporated with the target plasmid into the *E. coli* MG1655 cells harboring pORTMAGE-Ec1.

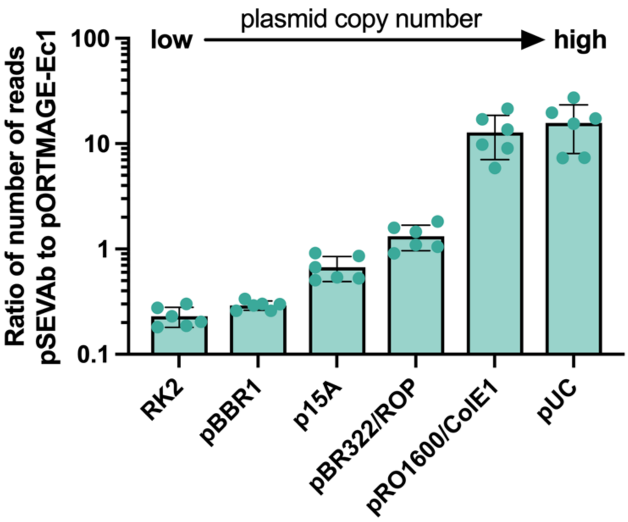


**Figure S4. Ratio of the number of Nanopore sequencing reads for the pSEVAb plasmids to those for pORTMAGE-Ec1.** *E. coli* MG1655 cells harboring pORTMAGE-Ec1 were electroporated with one of the pSEVAb plasmids and a mutagenic oligo. After plasmid isolation and Nanopore sequencing, the number of sequencing reads corresponding to the pSEVAb or pORTMAGE-Ec1 plasmids were determined by counting the reads with a size corresponding to that of the plasmids (± 100 bp). Then, the ratio of pSEVAb reads to pORTMAGE-Ec1 reads was calculated. pSEVAb plasmids with higher copy number origins of replication exhibited correspondingly elevated pSEVAb-to-pORTMAGE-Ec1 read ratios, thus affirming the consistency of the copy numbers reported in literature with the experimental (relative) copy numbers in our study (1, 2).

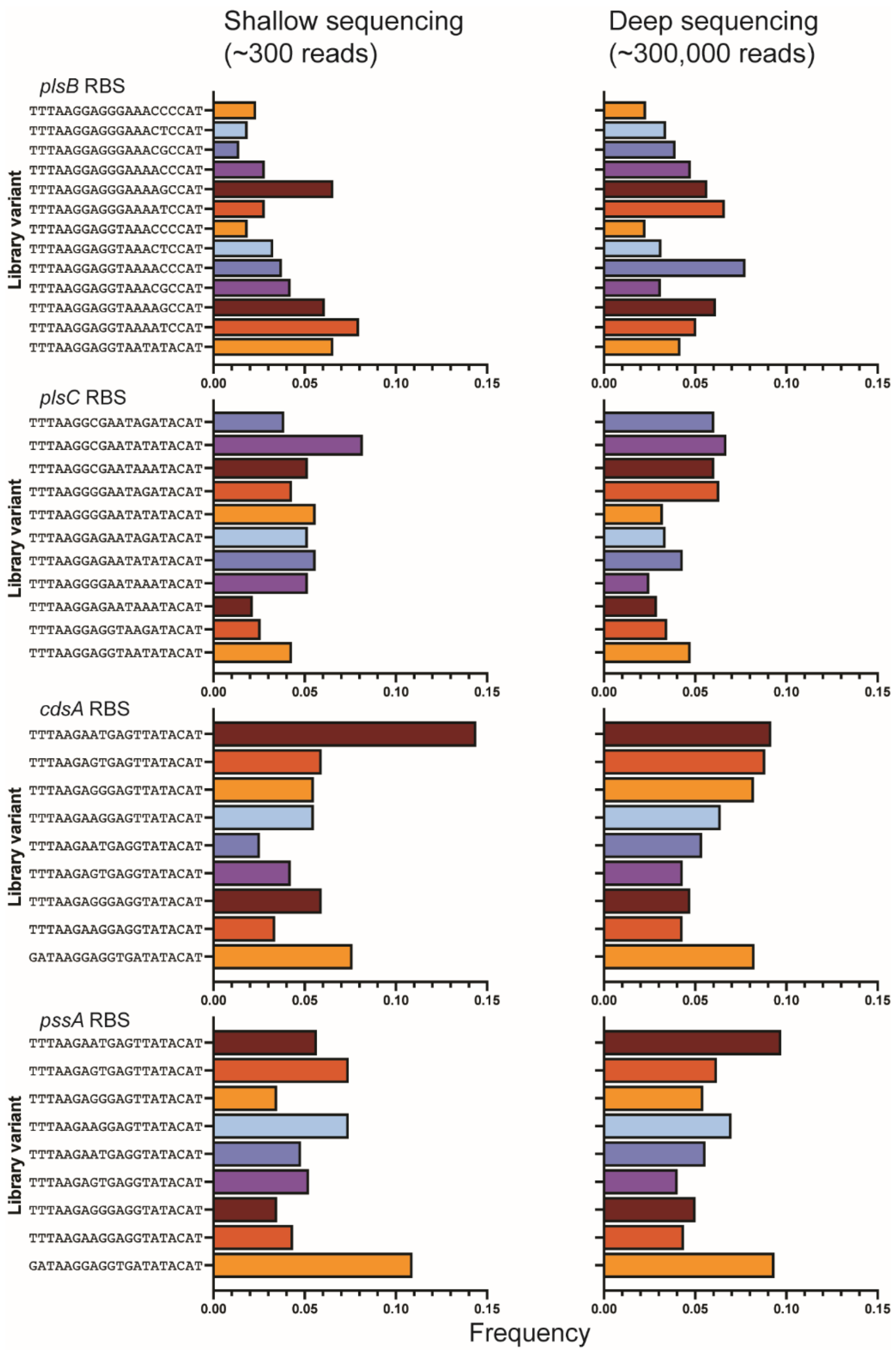


**Figure S5. Frequency of each of the RBS variants per target locus.** The graphs display the data for one of the four replicates from Figure 2C. The variant frequencies were quantified from shallow Nanopore sequencing (n=4, 150-300 reads) or deep sequencing (n=1, 279,454 reads).

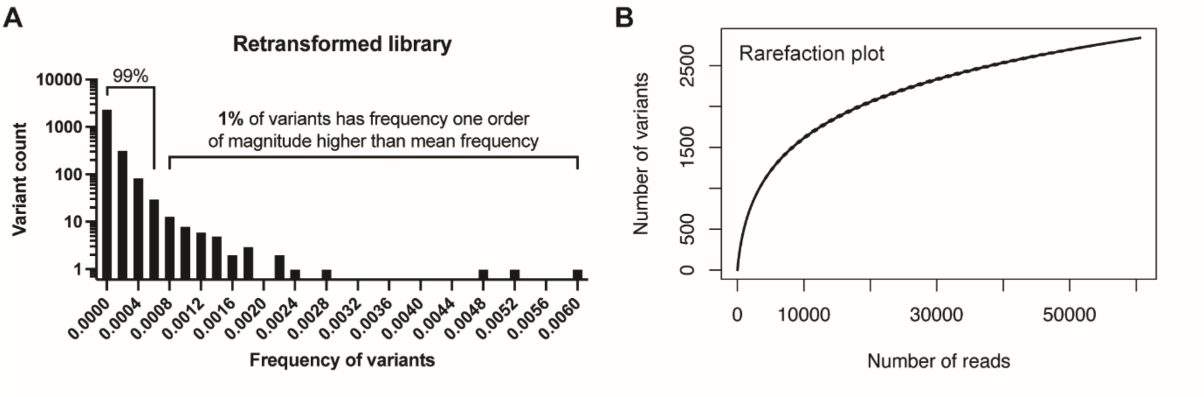


**Figure S6. Sequencing depth for the retransformed RBS library. A)** Frequency distribution of designed library variants in the deep sequencing dataset (279,454 reads). Sequencing reads containing at least one unmodified RBS were not considered. The average frequency was 0.0001 ± 0.0002 (mean ± standard deviation). **B)** Rarefaction plot of the sequenced DNA variants with all four RBSs mutated.

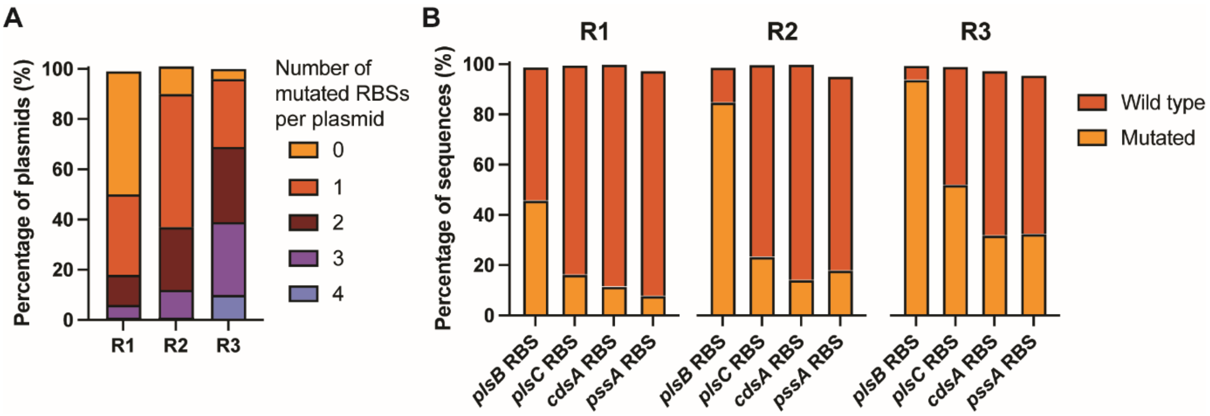


**Figure S7. Three successive rounds of MOSAIC increase the RBS library diversity of plasmid G555.** Strain used: DH5α. Oligos used: *1544 ChD*, *1545 ChD* and *1549 ChD* (Table S1). **A)** Number of mutated RBSs per plasmid. **B)** Percentage of wild-type and library sequences per target locus. The stacked bars do not match up to exactly 100%, because a small fraction of the sequences was neither the wild-type sequence nor a designed library variant. These unintended mutations were likely sequencing errors that were not excluded by our analysis pipeline (Figure S2B), mutations incorporated by oligos intended to bind at other target sites, and/or spontaneous mutations retained due to the suppression of the mismatch repair system during recombineering.

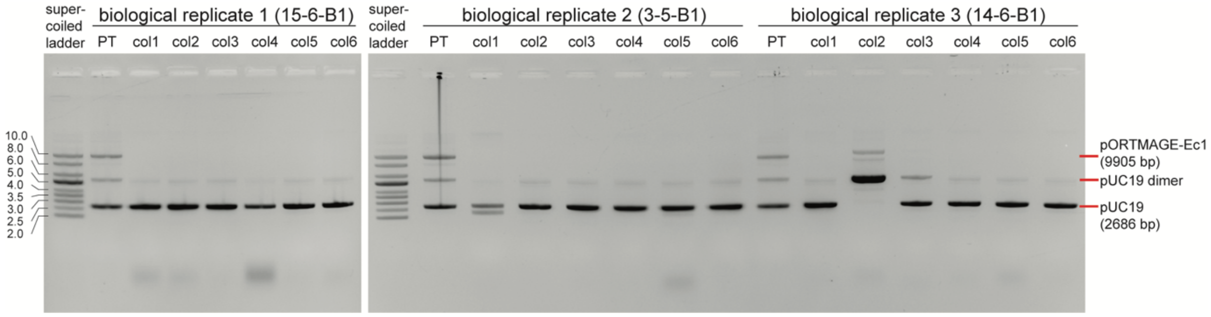


**Figure S8. Isolation of mutated pUC19 by retransformation.** pUC19 was mutated in the *lacZ* gene as shown in Figure 1B (3 mismatches, co-electroporation, MG1655). The resulting plasmid mixtures (n=3 biological replicates), containing mutated pUC19, wild-type pUC19 and pORTMAGE-Ec1, were used for chemical transformation of *E. coli* DH5α cells. Per sample, 6 colonies were picked from selective agar plates, grown in liquid LB medium, and plasmids were isolated from each cell culture. 100 ng of plasmids were run on a 0.7% agarose gel. Plasmid sizes were determined using the supercoiled DNA ladder (NEB) with plasmids ranging from 2.0-10.0 kb. PT=pre-transformation plasmid mixture containing pUC19 and pORTMAGE-Ec1. Col[x]=isolated plasmids from colony [x] after transformation. Nanopore sequencing results of some of the isolated plasmids are presented in Figure 3B. A small fraction of pUC19 dimer was formed, as indicated in the figure. The formation of multimers during plasmid recombineering has been reported before (3, 4). However, in our experiment, it remained unclear whether the dimers were formed due to the expressed recombineering machinery, or due to an alternative mechanism in *E. coli* MG1655 and/or in DH5α. Due to the small fraction of dimers formed as compared to the monomer, this did not cause a problem during the isolation of mutated pUC19 monomer.

**Table S1. DNA oligos used in this study.** Phosphorothioate bonds are indicated by an asterisk (*). Degenerate oligos (DEG) targeting G555 were mixed with single oligos (SINGLE) as additional library variants before mixing of oligos, plasmid and cell suspension for electroporation.

| Oligo name | Sequence | Target DNA | Description of use in this study |
|---|---|---|---|
| 1285 ChD / 1286 ChD | AACCATGGCTAGAACGTCTCAATCTCTATTAATAC / TTCTCGAGTTACAGGATGCGGCTAATTAATCGGTC | Plasmid G363 | Forward and reverse PCR primers for amplification of fragment containing four genes of phospholipid synthesis pathway for the assembly of plasmid G555 |
| 1287 ChD / 1288 ChD | AACTCGAGTAGCATAACCCCTTGGGGCCTCTAAAC / AACCATGGACATGCGACACAGACGAAGCGCTAAAAC | Plasmid G340 | Forward and reverse PCR primers for amplification of backbone for the assembly of plasmid G555 |
| 1495 ChD | C*T*GGCGAAAGGGGGATGTGCTGCAAGGCGATTAAGTTGGGTAACGCAGGGTTTTCCCAGTCACGACGTTGTAAAACGACGGCCAGTGAAT | pUC19/ *E. coli* genome | Single deletion in *lacZ* gene (introducing frame shift), lagging strand oligo for pUC19, leading strand oligo for *E. coli* genome |
| 1496 ChD | C*T*GGCGAAAGGGGGATGTGCTGCAAGGCGATTAAGTTGGGTAGCTCAAGGGTTTTCCCAGTCACGACGTTGTAAAACGACGGCCAGTGAAT | pUC19/ *E. coli* genome | 3-nt mismatch in *lacZ* gene (introducing stop codon), lagging strand oligo for pUC19, leading strand oligo for *E. coli* genome |
| 1489 ChD | A*T*TCACTGGCCGTCGTTTTACAACGTCGTGACTGGGAAAACCCTGCGTTACCCAACTTAATCGCCTTGCAGCACATCCCCCTTTCGCCAG | pUC19/ *E. coli* genome | Single deletion in *lacZ* gene (introducing frame shift), leading strand oligo for pUC19, lagging strand oligo for *E. coli* genome |
| 1490 ChD | A*T*TCACTGGCCGTCGTTTTACAACGTCGTGACTGGGAAAACCCTTGAGCTACCCAACTTAATCGCCTTGCAGCACATCCCCCTTTCGCCAG | pUC19/ *E. coli* genome | 3-nt mismatch in *lacZ* gene (introducing stop codon), leading strand oligo for pUC19, lagging strand oligo for *E. coli* genome |
| 1505 ChD | C*T*GGCGAAAGGGGGATGTGCTGCAAGGCGATTAAGTGTCCGCTGAAGGAATCGATCCCAGTCACGACGTTGTAAAACGACGGCCAGTGAAT | pUC19 | 18-nt mismatch in *lacZ* gene, lagging strand oligo for pUC19 |
| 1506 ChD | T*T*ACGCCAGCTGGCGAAAGGGGGATGTGCTGCAAGGCGATTAAGTTCCCAGTCACGACGTTGTAAAACGACGGCCAGTGAATTCGAGCTCG | pUC19 | 18-nt deletion in *lacZ* gene, lagging strand oligo for pUC19 |
| 1507 ChD | A*G*GGGGATGTGCTGCAAGGCGATTAAGTTGGGTAACAAAGTATCAGGACGGTATGCCAGGGTTTTCCCAGTCACGACGTTGTAAAACGACG | pUC19 | 18-nt insertion in *lacZ* gene, lagging strand oligo for pUC19 |
| BG31270 | G*T*CATAAGTTTAGCGTTCGCGGTGAAGGTGAGGGCGACGCGACCACGGCAAACTGACCCTGAAGTTCATCTGCACCACCGGTAAACTGC | pSEVAb plasmids | Single deletion in *sfGFP* gene, leading strand oligo |

| | | | |
|---|---|---|---|
| BG31271 | G*T*CATAAGTTTAGCGTTCGCGGTGAAGGTGAGGGCGACGCGACCACCAGTAAACTGACCCTGAAGTTCATCTGCACCACCGGTAAACTGC | pSEVAb plasmids | 3-nt mismatch in *sfGFP* gene, leading strand oligo |
| BG31272 | G*C*AGTTTACCGGTGGTGCAGATGAACTTCAGGGTCAGTTTGCCGTGGTCGCGTCGCCCTCACCTTCACCGCGAACGCTAAACTTATGAC | pSEVAb plasmids | Single deletion in *sfGFP* gene, lagging strand oligo |
| BG31273 | G*C*AGTTTACCGGTGGTGCAGATGAACTTCAGGGTCAGTTTACTGGTGGTCGCGTCGCCCTCACCTTCACCGCGAACGCTAAACTTATGAC | pSEVAb plasmids | 3-nt mismatch in *sfGFP* gene, lagging strand oligo |
| PlsB DEG | C*A*ACGGTTTCCCTCTAGAAATAATTTTGTTTAACTTTAAGGAGGKAAAMBCCATATGACTTTCTGCTATCCTTGCCGCGCATTTGCATTA | Plasmid G555 | 12-variant library targeting RBS of *plsB* gene, lagging strand oligo |
| PlsC DEG | C*A*ACGGTTTCCCTCTAGAAATAATTTTGTTTAACTTTAAGGVGAATADATACATATGCTATATATCTTTCGTCTTATTATTACCGTGATT | Plasmid G555 | 9-variant library targeting RBS of *plsC* gene, lagging strand oligo |
| PssA_CdsA DEG | A*T*AACAATTCCCCTCTAGAAATAATTTTGTTTAACTTTAAGARKGAGKTATACATATGGCTAGCATGACTGGTGGACAGCAAATGGGTCG | Plasmid G555 | 8-variant library targeting RBSs of *cdsA* and *pssA* genes, lagging strand oligo |
| PlsB SINGLE | C*A*ACGGTTTCCCTCTAGAAATAATTTTGTTTAACTTTAAGGAGGTAATATACATATGACTTTCTGCTATCCTTGCCGCGCATTTGCATTA | Plasmid G555 | Single library variant targeting RBS of *plsB* gene, mixed with degenerate oligo set before recombineering, lagging strand oligo |
| PlsC_1 SINGLE | C*A*ACGGTTTCCCTCTAGAAATAATTTTGTTTAACTTTAAGGAGGTAATATACATATGCTATATATCTTTCGTCTTATTATTACCGTGATT | Plasmid G555 | Single library variant targeting RBS of *plsC* gene, mixed with degenerate oligo set before recombineering, lagging strand oligo |
| PlsC_2 SINGLE | C*A*ACGGTTTCCCTCTAGAAATAATTTTGTTTAACTTTAAGGAGGTAAGATACATATGCTATATATCTTTCGTCTTATTATTACCGTGATT | Plasmid G555 | Single library variant targeting RBS of *plsC* gene, mixed with degenerate oligo set before recombineering, lagging strand oligo |
| PssA_CdsA SINGLE | A*G*CGGATAACAATTCCCCTCTAGAAATAATTTTGTTTAACGATAAGGAGGTGATATACATATGGCTAGCATGACTGGTGGACAGCAAATG | Plasmid G555 | Single library variant targeting RBSs of *cdsA* and *pssA* genes, mixed with degenerate oligo set before recombineering, lagging strand oligo |
| 1544 ChD | C*A*ACGGTTTCCCTCTAGAAATAATTTTGTTTAACTTTAAGGAGGGAAAACCCATATGACTTTCTGCTATCCTTGCCGCGCATTTGCATTA | Plasmid G555 | Single library variant targeting RBS of *plsB* gene, lagging strand oligo *Not used to generate RBS library, used only for Figure S7* |
| 1545 ChD | C*A*ACGGTTTCCCTCTAGAAATAATTTTGTTTAACTTTAAGGAGAATATATACATATGCTATATATCTTTCGTCTTATTATTACCGTGATT | Plasmid G555 | Single library variant targeting RBS of *plsC* gene, lagging strand oligo *Not used to generate RBS library, used only for Figure S7* |

| | | | |
|---|---|---|---|
| 1549 ChD | A*T*AACAATTCCCCTCTAGAAAT AATTTTGTTTAACTTTAAGAATGA GTTATACATATGGCTAGCATGAC TGGTGGACAGCAAATGGGTCG | Plasmid G555 | Single library variant RBSs of *cdsA* and *pssA* genes, lagging strand oligo *Not used to generate RBS library, used only for Figure S7* |

**Table S2. Number of sequenced DNA variants per colony.** From the Nanopore reads that span all four target loci, the different DNA variants were counted. The average number of DNA variants per colony was 6 ± 4 (n=6 colonies).

| Colony # | DNA variants (shown are the four target sequences: PlsB RBS … PlsC RBS … CdsA RBS … PssA RBS) | Count |
|---|---|---|
| 1 (total 60 reads, 5 different DNA variants) | TTTAAGGAGGTAAACCCCAT…TTTAAGAAGGAGATATACAT… TTTAAGAAGGAGATATACAT…TTTAAGAGTGAGGTATACAT | 43 |
| | TTTAAGGAGGAGATATACAT…TTTAAGGAGGAGATATACAT… TTTAAGAAGGAGTTATACAT…TTTAAGAAGGAGATATACAT | 8 |
| | TTTAAGAAGGAGATATACAT…TTTAAGAAGGAGATATACAT… TTTAAGAAGGAGATATACAT…TTTAAGAAGGAGATATACAT | 7 |
| | TTTAAGGAGGTAAAATCCAT…TTTAAGGAGGAGATATACAT… TTTAAGAAGGAGTTATACAT…TTTAAGAAGGAGATATACAT | 1 |
| | TTTAAGGAGGTAAACCCCAT…TTTAAGAAGGAGATATACAT… TTTAAGAAGGAGATATACAT…TTTAAGAAGGAGATATACAT | 1 |
| 2 (total 22 reads, 11 different DNA variants) | TTTAAGAAGGAGATATACAT…TTTAAGGGGAATAGATACAT… TTTAAGAAGGAGTTATACAT…TTTAAGAAGGAGATATACAT | 7 |
| | TTTAAGAAGGAGATATACAT…TTTAAGGGGAATAGATACAT… TTTAAGAAGGAGATATACAT…TTTAAGAAGGAGATATACAT | 1 |
| | TTTAAGAAGGAGATATACAT…TTTAAGAAGGAGATATACAT… TTTAAGAAGGAGTTATACAT…GATAAGGAGGTGATATACAT | 1 |
| | TTTAAGAAGGAGATATACAT…TTTAAGAAGGAGATATACAT… TTTAAGAAGGAGATATACAT…GATAAGGAGGTGATATACAT | 1 |
| | TTTAAGAAGGAGATATACAT…TTTAAGGGGAATAGATACAT… TTTAAGAAGGAGTTATACAT…GATAAGGAGGTGATATACAT | 1 |
| | TTTAAGAAGGAGATATACAT…TTTAAGAAGGAGATATACAT… TTTAAGAAGGAGATATACAT…TTTAAGAAGGAGATATACAT | 4 |
| | TTTAAGAAGGAGATATACAT…TTTAAGAAGGAGATATACAT… TTTAAGAAGGAGTTATACAT…TTTAAGAAGGAGATATACAT | 1 |
| | TTTAAGAAGGAGATATACAT…TTTAAGAAGGAGATATACAT… TTTAAGAAGGAGGTATACAT…TTTAAGAAGGAGATATACAT | 2 |
| | TTTAAGAAGGAGATATACAT…TTTAAGAAGGAGATATACAT… TTTAAGAAGGAGGTATACAT…GATAAGGAGGTGATATACAT | 1 |
| | TTTAAGAAGGAGATATACAT…TTTAAGGAGAATATATACAT…T TTAAGAAGGAGTTATACAT…GATAAGGAGGTGATATACAT | 2 |
| | TTTAAGAAGGAGATATACAT…TTTAAGGAGAATATATACAT…T TTAAGAAGGAGGTATACAT…TTTAAGAAGGAGATATACAT | 1 |
| 3 (total 30 reads, 9 different DNA variants) | TTTAAGAAGGAGATATACAT…TTTAAGGCGAATATATACAT…T TTAAGAGGGAGGTATACAT…TTTAAGAAGGAGGTATACAT | 2 |
| | TTTAAGGAGGTAAACGCCAT…TTTAAGAAGGAGATATACAT… TTTAAGAAGGAGATATACAT…TTTAAGAAGGAGATATACAT | 1 |
| | TTTAAGGAGGTAAAATCCAT…TTTAAGAAGGAGATATACAT… TTTAAGAAGGAGATATACAT…TTTAAGAAGGAGATATACAT | 4 |

| | | |
|---|---|---|
| | TTTAAGAAGGAGATATACAT...TTTAAGGAGGTAAGATACAT...<br>TTTAAGAAGGAGGTATACAT...TTTAAGAAGGAGATATACAT | 3 |
| | TTTAAGAAGGAGATATACAT...TTTAAGAAGGAGATATACAT...<br>TTTAAGAGGGAGGTATACAT...TTTAAGAAGGAGATATACAT | 11 |
| | TTTAAGAAGGAGATATACAT...TTTAAGAAGGAGATATACAT...<br>TTTAAGAAGGAGATATACAT...TTTAAGAAGGAGATATACAT | 3 |
| | TTTAAGAAGGAGATATACAT...TTTAAGAAGGAGATATACAT...<br>TTTAAGAGGGAGGTATACAT...GATAAGGAGGTGATATACAT | 1 |
| | TTTAAGGAGGTAAAACCCAT...TTTAAGAAGGAGATATACAT...<br>GATAAGGAGGTGATATACAT...TTTAAGAAGGAGATATACAT | 2 |
| | TTTAAGAAGGAGATATACAT...TTTAAGGAGAATATATACAT...T<br>TTAAGAAGGAGATATACAT...TTTAAGAAGGAGATATACAT | 3 |
| 4<br>(total 29 reads,<br>4 different DNA variants) | TTTAAGAAGGAGATATACAT...TTTAAGAAGGAGATATACAT...<br>TTTAAGAGTGAGGTATACAT...TTTAAGAAGGAGATATACAT | 10 |
| | TTTAAGGAGGGAAACCCCAT...TTTAAGAAGGAGATATACAT...<br>TTTAAGAAGGAGATATACAT...TTTAAGAAGGAGATATACAT | 5 |
| | TTTAAGGAGGGAAAATCCAT...TTTAAGAAGGAGATATACAT...<br>TTTAAGAATGAGTTATACAT...TTTAAGAGTGAGTTATACAT | 13 |
| | TTTAAGAAGGAGATATACAT...TTTAAGAAGGAGATATACAT...<br>TTTAAGATGAAGGTATACAT...TTTAAGAAGGAGATATACAT | 1 |
| 5<br>(total 27 reads,<br>3 different DNA variants) | TTTAAGAAGGAGATATACAT...TTTAAGAAGGAGATATACAT...<br>TTTAAGAAGGAGATATACAT...TTTAAGAAGGAGATATACAT | 23 |
| | TTTAAGAAGGAGATATACAT...TTTAAGAAGGAGATATACAT...<br>TTTAAGAGGGAGGTATACAT...TTTAAGAAGGAGATATACAT | 1 |
| | TTTAAGAAGGAGATATACAT...TTTAAGAAGGAGATATACAT...<br>TTTAAGAATGAGGTATACAT...TTTAAGAAGGAGATATACAT | 3 |
| 6<br>(total 29 reads,<br>2 different DNA variants) | TTTAAGAAGGAGATATACAT...TTTAAGAAGGAGATATACAT...<br>TTTAAGAAGGAGATATACAT...TTTAAGAAGGAGATATACAT | 28 |
| | TTTAAGAAGGAGATATACAT...TTTAAGGAGGTAAGATACAT...<br>TTTAAGAAGGAGATATACAT...TTTAAGAAGGAGATATACAT | 1 |
